# Tolerance to land-use changes through natural modulations of the plant microbiome

**DOI:** 10.1101/2023.11.13.566857

**Authors:** Vincent Zieschank, Anne Muola, Robert R. Junker

**Affiliations:** Evolutionary Ecology of Plants, Department of Biology, Philipps-University Marburg, 35043 Marburg, Germany; Division of Biotechnology and Plant Health, Norwegian Institute of Bioeconomy Research, 9016 Tromsø, Norway

**Keywords:** bacteria, fertilizer, fungi, microbiome, mowing, plant performance, plant phenotype

## Abstract

Land-use changes pose a threat to many ecosystems and are a major driver of species loss. Adaptations to altered environments or migration to more suitable habitats are potential mechanisms to resist global change, which can, however, lag behind rapid anthropogenic alterations of the environment. Our data show that rapid natural modulations of the plant microbiome in response to land-use change directly affect plant phenotype and performance, and thus increase plant tolerance to environmental changes. In a common garden experiment, the effects of the microbiome on the plant phenotype were stronger than the direct effects of fertilizer application and mowing. This finding was confirmed in a subsequent controlled laboratory experiment using plants inoculated with land-use-specific microbiomes. Therefore, natural modulations of the plant microbiome may be the key to species persistence and ecosystem stability. A prerequisite for this microbiome-mediated tolerance is the availability of diverse local sources of microorganisms that can function as a resource for rapid modulations in response to change. Thus, conservation efforts must protect microbial diversity, which can help mitigate the effects of global change and facilitate environmental and human health.

## Introduction

Land-use changes are a major driver of species loss and alterations in community structure with consequences for ecosystem functioning and human well-being (Allan et al., 2015; Díaz et al., 2019). Plant species are increasingly challenged to cope with these environmental alterations. Migration to more suitable habitats or adaptations to novel conditions are frequently discussed mechanisms to avoid extinction (Feeley et al., 2012; Scheffers et al., 2016), which may, however, lag behind rapid anthropogenic alterations of environmental conditions (Etterson & Shaw, 2001). Natural modulations of plant-associated microbial communities in response to environmental stresses can represent an alternative – and faster – way for plants to cope with alterations in land-use and other components of global change (Afkhami, 2023; Allsup et al., 2023; Lau & Lennon, 2012; Trivedi et al., 2022). In fact, ecological and evolutionary microbiome responses can easily keep pace with environmental changes (Martiny et al., 2023) and land-use history, intensity, and agricultural practices have been shown to affect the diversity and composition of bacterial and fungal communities in the soil or associated with plants (Abrahao et al., 2022; Bai et al., 2023; Estendorfer et al., 2017; Gaube et al., 2021; Li et al., 2019). These rapid responses of plant-associated microbiomes may then, in turn, have positive effects on plant phenotype and performance, increasing plant species’ tolerance to changes (Friesen et al., 2011; Jiang et al., 2023). As a consequence, plants may invariantly perform well despite being challenged by stressors (Kiesewetter & Afkhami, 2021; Porter et al., 2020; Timmusk et al., 2014). Accordingly, inoculation experiments in the context of climate change showed that a preadapted microbiome can increase survival rates of seedlings and promote plant growth under environmental stress (Allsup et al., 2023; Carrell et al., 2022; Timmusk et al., 2014). Positive microbial effects on plant species could even promote species richness and productivity in plant communities (Abrahao et al., 2022; Laforest-Lapointe et al., 2017; Li et al., 2019). Therefore, spontaneous natural changes in plant-associated microbiomes may be equivalent to recent advances in microbiome engineering, which is used to promote plant fitness and support plant tolerance to rapid changes of the environment (Rodriguez & Duran, 2020).

Especially in grasslands, land degradation and changes in land-use intensity have been identified as the main causes of diversity decline, disruptions in species interactions, and loss of ecological functions (Allan et al., 2015; Gossner et al., 2016; Manning et al., 2015; Newbold et al., 2015). Different components of land-use (e.g. mowing or fertilizer application) have been shown to affect plant and microbial diversity and composition differently (Gaube et al., 2021; Kaiser et al., 2016; Socher et al., 2013; Zieschank & Junker, 2023). Although some taxa benefit from an increase in land-use intensity, many decline and are eventually displaced from their original habitat (Allan et al., 2015). Plants in natural and agricultural systems may benefit from microbes helping them resist abiotic stresses or protecting them against diseases and herbivores (Carrell et al., 2022; Kim et al., 2011; Timmusk et al., 2014; Yang et al., 2009). Both individual strains of microbes as well as diverse consortia can affect plant traits, plant interactions, and ecosystem processes and functions (Finkel et al., 2017; Grigulis et al., 2013; Junker et al., 2021; Laforest-Lapointe et al., 2017), but inoculated consortia usually outperform individual strains (Liu et al., 2023).

Based on the observed vulnerability of grassland species and the responses of bacteria and fungi to land-use changes, we tested whether natural rapid modulations of the plant microbiome mediate land-use effects on plant phenotype and performance and thus represent an alternative to ‘adapt or migrate’. In a common garden experiment, grass sods from different regions in Germany complemented with *Fragaria vesca* plants as phytometers were subjected to different mowing and fertilizer regimes (Zieschank & Junker, 2023) to test the effects of land-use treatments on morphological and physiological features of plant communities, *F. vesca* phenotype and performance, as well as the diversity and composition of the bacterial and fungal communities associated with *F. vesca*. Such experiments may reveal strong microbial contributions to land-use effects on plant phenotype and performance (Allsup et al., 2023; Carrell et al., 2022; Laforest-Lapointe et al., 2017), but clear evidence for direct effects of the plant microbiome modulated by global change on plant responses requires additional, more controlled laboratory experiments. Therefore, we inoculated *F. vesca* plants grown from surface sterilized seeds in germ-free containers with the whole microbiome extracted from field-grown phytometers, testing microbial effects on the plant phenotype under gnotobiotic conditions in the absence of environmental variation. In these common garden and laboratory experiments, we tested a set of hypotheses specifically addressing the prerequisites underlying the hypotheses of microbiome-assisted plant tolerance to land-use changes: 1) microbial diversity and composition as well as single plant phenotype and plant community features rapidly respond to major components of land-use, mowing and fertilization; 2) microbes affect plant phenotype and performance as strong or even stronger than direct land-use effects; 3) plant performance is invariant across land-use treatments; 4) microbial effects are also measurable in the absence of other biotic or abiotic influences. By verifying these hypotheses, we show that rapid natural modulations of the plant microbiome increase plant tolerance to land-use changes and therefore may also buffer grasslands against stressors of global change.

## Material and methods

### Plant communities

The project is part of the Biodiversity Exploratories, a large-scale research network with three long-term research sites (‘Exploratories’) in Germany at Schwäbische Alb, Hainich, and Schorfheide-Chorin, where land-use and biodiversity of 50 grassland plots per region have been continuously recorded (Fischer et al., 2010). In 2020, we established a common-garden (‘EXClAvE’) to track effects of land-use by reducing environmental heterogeneity to a minimum. In each region, we selected 13 plots that evenly cover the range of land-use intensity (measured as land-use index LUI, see Blüthgen et al., 2012). On every pre-selected plot, we collected grass sods the size of 1m^2^ and 10cm depth in April and May 2020. The sods were split into four parts of 50×50cm and each part was subsequently randomly assigned to one of four experimental land-use treatments: ‘00’ (= mowing once per year), ‘0M’ (= mowing twice per year), ‘0F’ (= mowing once per year and fertilizer addition) and ‘MF’ (= mowing twice per year and fertilizing). Together with 12 sods made from local steam-sterilized soil to monitor spontaneous plant establishment from the surroundings, the common garden consists of 168 sods/plant communities, (for more information see Zieschank & Junker, 2023). Land-use treatments first started in mid-July 2020. Plant communities were mown in early summer and autumn each year and fertilization (99 kg N ha^-1^ year^-1^) was done right after the first mowing, respectively (see Zieschank & Junker, 2023).

### Digital phenotyping

For data collection, we customized an automated plant phenotyping system (PlantEye F500, Phenospex, Heerlen, The Netherlands) for its mobile application in the field (described in Zieschank & Junker, 2023). By scanning whole plant communities, we gather, within seconds and non-invasively, multispectral and physiological information while simultaneously capturing the 3-dimensional structure of the vegetation. The scans are processed with the built-in software HortControl (Phenospex, Heerlen, The Netherlands) that provides multiple morphological and physiological parameters (Zieschank & Junker, 2023). The accuracy of this method in tracking plant community responses to mowing and fertilization was demonstrated in (Zieschank & Junker, 2023). For this study we excluded the parameter ‘maximum height’ from all analyses because of its unsuitability for scans of whole communities. The scans of plant communities used in this study were recorded in August 2021.

### *Fragaria* field experiment

We used *Fragaria vesca* plants (accession: *PI616612, USDA National Clonal Germplasm Repository, Corvallis, US)* as phytometers in the common garden because this species has been recorded at the grassland plots of the Biodiversity Exploratories but does not naturally occur on any of the chosen plots sampled for the common garden. *F. vesca* is an herbaceous perennial plant occurring throughout the Northern Hemisphere(Hilmarsson et al., 2017) that grows in various habitats such as grasslands, roadsides, open forests, and forest and farmland edges (Roiloa & Retuerto, 2007; Schulze et al., 2012). *F. vesca* flowers are hermaphrodite and self-compatible to various degrees(Angevine, 1983; Egan et al., 2018). *F. vesca* is insect pollinated and reproducing both sexually and clonally through formation of aboveground runners (Muola et al., 2017; Schulze et al., 2012). To prepare the experiment, we grew *n* = 380 individuals of a clone of *F. vesca* in a greenhouse. Based on morphological and physiological parameters from digital phenotyping (Zieschank & Junker, 2023), we selected a subset of *n* = 168 plants with a homogenous phenotype. Immediately prior to transferring the plants to the common garden, all *F. vesca* plants were digitally phenotyped. Each plant was scanned from two opposite sides, which were marked on the pot for repeated scans at the end of the experiment. The mean value of the two scans was used as dependent variable. We planted one individual of *F. vesca* into each sod of the common garden in May 2021. The plants remained in their pots with the original soil in order to exclude soil and nutrient effects of the sods on plant growth and performance. After a one-week acclimatization period, *F. vesca* plants received the same land-use treatment as the surrounding plant community: ‘00’ (= no treatment), ‘M0’ (= mowing), ‘0F’ (= fertilizer addition) and ‘MF’ (= mowing and fertilizing). For the mowing treatment, plants were cut at about 2cm above ground and the fertilizer quantity was adjusted to match the fertilization of the sods (99 kg N ha^-1^ year^-1^). After three months, we recorded herbivory (visual assessment of % of leaf damage after online training using the ZAX Herbivory TrainerXirocostas et al., 2022), fruit set (number of fruits and flowers) and vegetative reproduction (number of stolons and offshoots) as proxies for performance. *Fragaria vesca* plants were scanned again from the previously marked sides and one leaf and root sample was taken from each plant for next generation sequencing. To account for small differences in the phyotmeters’ phenotype prior to the experiment, we used the differences in phenotypic parameters between the scan after and before the experiment as dependent variable in all further statistical analyses.

### Microbiome analysis

Leaf and root samples of *F. vesca* phyotmeters were collected using sterilized forceps (dipped into 70% ethanol and flamed) to avoid contamination. Samples were directly transferred to ZR BashingBead Lysis tubes containing 750μL of ZymoBIOMICS lysis solution (Zymo-BIOMICS DNA Miniprep Kit, Zymo Research, Irvine, CA). ZR BashingBead Lysis tubes were sonicated for 7 minutes right after the collection of microbial samples to detach microorganisms from the surfaces. After sonication, leaf and root tissue was removed from the tubes with sterile forceps to reduce the amount of plant DNA in the samples. Subsequently, all microbial samples were shaken using a ball mill for 9 minutes with a frequency of 30.0s–1. Microbial DNA was isolated using the ZymoBIOMICS DNA Miniprep Kit following the manufacturers’ instructions. Amplicon sequencing of isolated DNA samples was performed by Eurofins Genomics (Ebersberg, Germany). The V3-V4 region of the 16S rRNA and the ITS2 region of leaf and root microbiome samples were amplified and sequenced with Illumina MiSeq to identify bacterial and fungal communities, respectively. Microbiome profiling of isolated DNA was performed on the Qiita web platform (Gonzalez et al., 2018), where bacterial ASVs were obtained using Deblur (deblur 2021.09) (Amir et al., 2017) and fungal ASVs were obtained using QIIME 2 2021.8.0 (Bolyen et al., 2019). Taxonomy was assigned to bacterial ASVs using the q2-feature-classifier (Bokulich et al., 2018) classify-sklearn naïve Bayes taxonomy classifier against the Greengenes 13_8 99% OTUs reference sequences (McDonald et al., 2012). Raw sequences of the fungal data were demultiplexed and quality filtered using the q2-demux plugin followed by denoising with DADA2 (via q2-dada2) (Callahan et al., 2016). Taxonomy was assigned to fungal ASVs using the q2-feature-classifier classify-sklearn naïve Bayes taxonomy classifier against the UNITE 97% Version 8.3 reference sequences (Kõljalg et al., 2020). Prior to the statistical analysis of microbial communities, we performed a cumulative sum scaling (CSS) normalization (R package metagenomeSeq v1.28.2Paulson, 2014) on the count data to account for differences in sequencing depth among samples.

### *Fragaria* lab experiment

In order to transplant the full microbiome of the phytometers to individuals of the *F. vesca* clone, we first removed soil from roots and then placed the plants into plastic bags filled with 225 ml sterile water and shook them in a pulsifier (Microgen Bioproducts Ltd, Camberley, Surrey, UK) for 60 seconds (following the user manual). 1ml of each extract was stored in a freezer (−80°C) after adding 0.5ml Glycerin for future use in the follow-up inoculation experiment. Seeds from the same *F. vesca* clone were sterilized by sequential exposure to 70% ethanol (10min) and a sodium hypochlorite solution (5min) and four subsequent washes with sterile water (following the protocol of Engelsdorf et al., 2018). After 24 hours in the fridge, sterilized seeds were put individually into pre-autoclaved (121°C, 20min, 1.1bar) microcosm containers with tightly closing lids containing a filter that allows gas exchange but no passage of microorganisms (SacO2 O118/120+OD118/80, SacO2 NV, Veldeken 29, 9850 Deinze, Belgium). Each box was previously filled with 90g clay (Diamond Pro Calcined Clay drying agent, Diamond Pro, 1112 E. Copeland Rd, Ste 500, Arlington, TX 76011, Texas, USA) that has been washed repeatedly before drying and sterilizing it in a drying cabinet at 200°C for 24h. For nutrient and water supply we added autoclaved 50ml plant agar (5.5g L^-1^, Duchefa Biochemie B.V., 2003 RV Haarlem, The Netherlands) containing 1.1g L^-1^ Murashige & Skoog Medium including vitamins (Duchefa Biochemie B.V., 2003 RV Haarlem, The Netherlands) to each container. Two weeks after the emergence of the primary leaf, the plants were inoculated with the stored whole microbiome extract by pipetting each 10µl on three leaves of the plant. Per donor plant, two receiver plants were inoculated, resulting in *n* = 332 inoculated plants. Plants grew for three months in a climate chamber at 20°C and a 15h/9h light/dark cycle and were finally digitally phenotyped. Unfortunately, the scanner had an error during these scans, reducing the sample size from *n* = 166 to *n* = 95.

### Statistical analyses

We used structural equation modeling (i) to test the effect of mowing and fertilizing on plant communities, individual plant, the plant-associated microbiome diversity and composition, and plant performance; (ii) to investigate the role of the microbiome in mediating effects of mowing and fertilizing on plant phenotype and performance; and (iii) to compare the relative effect sizes of the microbiome and the land-use treatments on plant phenotype and performance. We built separate models with identical structure for microbiome diversity and microbiome composition. Exogenous variables in the model were mowing and fertilizing as categorical binary predictors (Y/N). Since the fixed range of categorical variables disrupts the presumed relationship between standard deviation and range of a predictor, thus interfering with the standardization of estimates within a model, we recalculated standardized estimates for relevant paths following the tutorial of Grace (2006), using defined-difference standardization where the defined difference is the known range of values(Grace & Bollen, 2005). Both models were estimated using the R-package *lavaan* (v0.6-12Rosseel, 2012).

For multivariate variables (plant community features, *Fragaria* phenotype, *Fragaria* performance and microbiome composition) in the SEM, we used principal components (PC1) as univariate representatives. PCA for plant community features and *Fragaria* phenotype included 13 parameters from digital phenotyping. *Fragaria* performance comprised inverse herbivory (% of undamaged leaf area), number of flowers, fruits, stolons and offshoots. For microbiome composition, we combined leaf and root ASV abundance data for bacteria and fungi, respectively, then merged bacterial and fungal ASVs for the final whole microbiome PCA. For microbiome diversity we first calculated Shannon diversities of bacteria and fungi from the combined leaf and root ASV abundance data (using non-normalized data). Second, we ranked samples by increasing diversity for bacteria and fungi individually and then calculated the mean rank per sample to get a cumulative microbiome diversity. For better interpretability we scaled mean ranks between zero and one. Ranks were used to give the same weighting to bacteria and fungi despite deviations in the absolute values of Shannon diversity(Hanusch et al., 2022). To account for the setup of the common garden, we tested for the presence of residual spatial autocorrelation in the estimated path models by calculating spatial neighbor matrices (k = 8) based on plant community coordinates and estimating Moran’s I for the case-wise residuals of the exogenous variables of both models using the package *spdep* (v1.2-7Bivand, 2022). Significant autocorrelation was only found for the residuals of the plant community variable in both models (*p* < 0.001, see Supplementary Table 9). Following a tutorial by Jim Grace (https://bit.ly/44kJ9gt, based on an approach from: Bivand et al. (2013)) we estimated the real effective sample size (n = 128), then adjusted standard errors accordingly and recalculated p-values for the model parameters affected by the autocorrelation.

We performed further analysis for a closer insight into the significant relationships between variables in the SEM. For a multivariate assessment of the effects of mowing and fertilizing on plant community, *Fragaria* phenotype, *Fragaria* performance and microbiome composition, we performed distance-based redundancy analyses using Bray-Curtis distances followed by a permutation test under reduced model with subsequent analysis of variance using functions *capscale* (R package vegan v2.6-2Oksanen et al., 2022) and *anova* (R base package stats 4.2.0R Core Team, 2022) with mowing and fertilizing separately as factors. Furthermore, we performed an *anova* (R base package stats 4.2.0R Core Team, 2022) with mowing and fertilizer as factors to explore the response of microbial diversity to land-use components and to look for treatment-associated microbiome-induced changes in the phenotype of previously inoculated lab plants. We used the default workflow steps implemented in the R package *DESeq2*, a tool for differential expression analysis, to trace significant changes triggered by mowing and fertilization in bacteria and fungi ASVs(Love et al., 2014). The strength of effects is measured as log2 fold change, which means the land-use component induced a multiplicative change of 2^-1^=0.5 in abundance of the observed ASV compared to plants that did not receive this treatment.

## Results

### Plants and microbiomes rapidly respond to land-use treatments

The plant communities of the transplanted grass sods that each contained one individual of a *Fragaria vesca* clone were subjected to one of four land-use treatments: ‘00’ (= mowing once per year), ‘0M’ (= mowing twice per year), ‘0F’ (= mowing once per year and fertilizer addition) and ‘MF’ (= mowing twice per year and fertilizing). Within the duration of the experiment, *F. vesca* plants received the following corresponding treatments: ‘00’ (= no treatment), ‘0M’ (= mowing), ‘0F’ (= fertilizer addition) and ‘MF’ (= mowing and fertilizing). Three months after *F. vesca* individuals were placed in the common garden, we assessed morphological and physiological features of the plant communities as well as *F. vesca* individuals by scanning them with a customized plant phenotyping system (PlantEye F500, Phenospex, Heerlen, The Netherlands). The plant scanner was originally constructed to scan plant individuals, but also provides meaningful information on whole plant communities (Zieschank & Junker, 2023). A total of *n* = 162 plant communities and *F. vesca* plants were included in the final analyses. In accordance with our first hypothesis, we found strong effects of mowing and fertilizer addition on plant communities (Constrained Analysis of Principal coordinates (CAP) based on Bray–Curtis distances followed by permutation test for capscale under reduced model; mowing: *F*_1,158_ = 7.87, *p* < 0.001; fertilizer: *F*_1,158_ = 7.27, *p* < 0.001; mowing x fertilizer: *F*_1,158_ = 1.96, *p* = 0.029; Fig. 1A).

**Figure 1.**
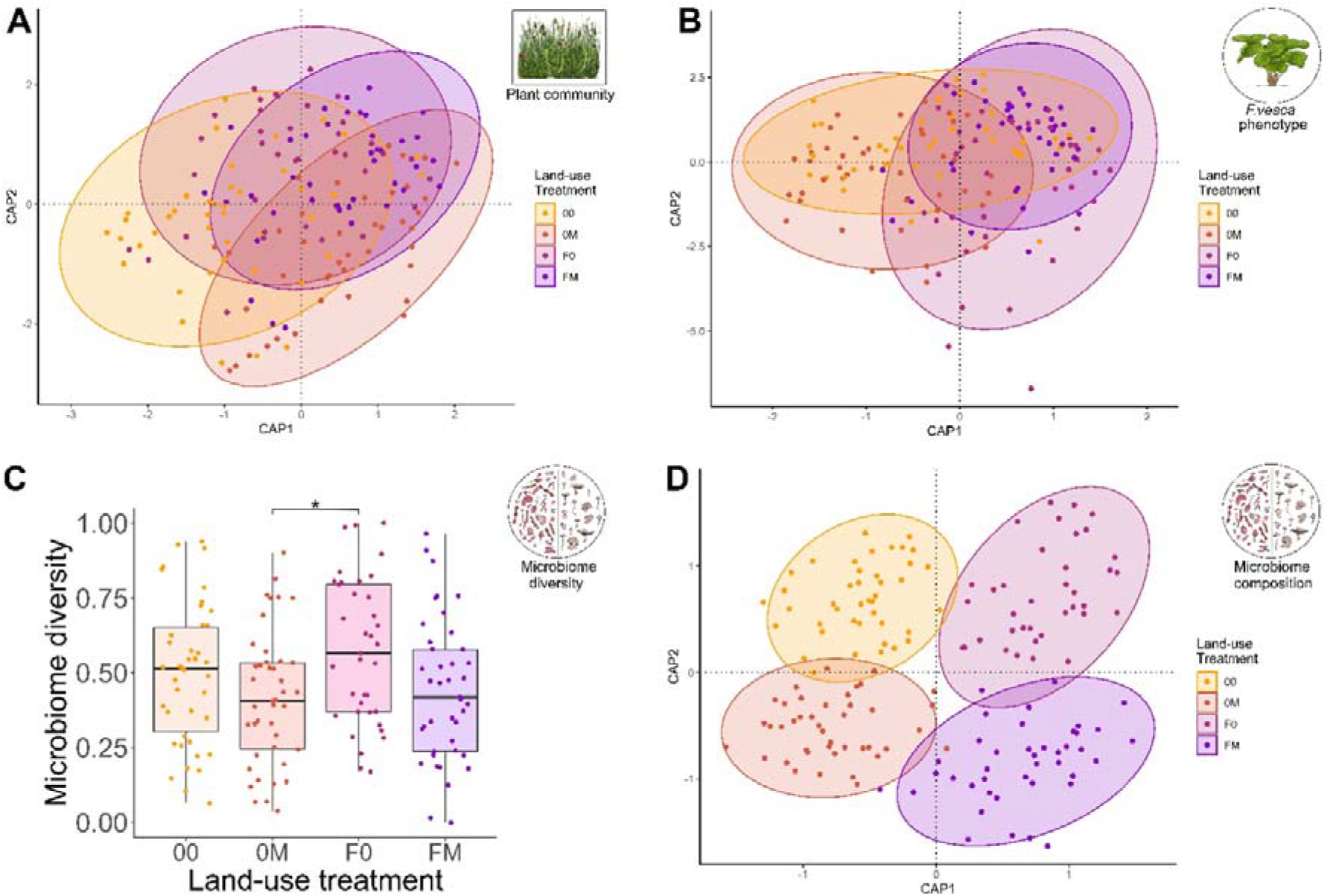
Land-use effects on plant communities, plant individuals and microbiomes in a common garden experiment. Plant community features (A) and the *F. vesca* phenotype (B) were assessed by 13 morphological and physiological parameters using a 3D plant scanner and both clearly responded to land-use treatments. Microbiome diversity (C) and composition (D) were analyzed based on 16S rRNA (bacteria) and ITS2 (fungi) amplicon sequencing and both, diversity and composition, responded to land-use treatments. Each circle represents one sample, land-use treatment is color-coded; ‘00’ (= mowing once per year), ‘0M’ (= mowing twice per year), ‘0F’ (= mowing once per year and fertilizer addition) and ‘MF’ (= mowing twice per year and fertilizing). Statistical results are reported in the text and Supplementary Material (Tables 1, 2, and 4).

Likewise, *F. vesca* plant individuals also clearly responded to land-use treatments by alterations in morphological and physiological features (fertilizer: *F*_1,158_ = 10.53, p < 0.001; mowing x fertilizer: *F*_1,158_ = 1.96, *p* = 0.029; Fig. 1B). Furthermore, bacterial and fungal communities associated with *F. vesca* roots and leaves (analyzed via next-generation 16S rRNA gene and ITS amplicon sequencing) strongly responded to land-use treatments, too. Microbial diversity responded negatively to mowing (ANOVA: *F*_1,158_ = 8.05, *p* = 0.005; Fig. 1C, see also Supplementary Table 1 for results on bacteria and fungi associated to leaves and roots separately) and land-use effects on microbiome composition were even more pronounced (CAP based on Bray–Curtis; mowing: F_1,158_ = 1.69, p < 0.001; fertilizer: F_1,158_ = 3.24, p < 0.001; Fig. 1D, see also Supplementary Table 2 for results on bacteria and fungi associated to leaves and roots separately). Next to effects on microbial communities, individual bacterial and fungal amplicon sequence variants (ASVs) responded to land-use treatments, too (Supplementary Figures 1 and 2). In total, we detected *n* = 4,486 bacterial ASVs and *n* = 791 fungal ASVs. The raw sequences of next-generation 16S and ITS amplicon sequencing are available at the European Nucleotide Archive ENA under the project accession PRJEB52951 (https://www.ebi.ac.uk/ena/browser/view/PRJEB52951).

### Effects of land-use and the plant-associated microbiome on plant phenotype and performance

We used structural equation modeling (SEM) to estimate the effect sizes of land-use components and plant microbiome on phenotype and performance of *F. vesca* plant individuals in a single directed analysis. For multivariate variables in the SEM, we used principal components (PC) as univariate representatives (morphological and physiological features of plant communities: PC1 = 45.37%; morphological and physiological features of *F. vesca* individuals: PC1 = 35.05%; performance of *F. vesca* individuals composed of herbivory, fruit set and flowers, vegetative reproduction: PC1 = 35.95%; microbiome composition: PC1 = 4.87%). We tested effects of microbial composition and diversity separately in two otherwise identical SEMs. Overall, we found a clear dependence structure between the variables included in the SEMs. *F. vesca* performance, which was invariant across land-use treatments (ANOVA; mowing: *F*_1,158_ = 1.98, *p* = 0.29; fertilizer: *F*_1,158_ = 2.51, *p* = 0.12), was solely affected by *F. vesca* phenotype in both SEMs (Fig. 2A, B), emphasizing the central role of the plants’ morphological and physiological traits. SEM results were supported by a clear correlation between phenotype and performance of *F. vesca* individuals (Mantel based on Pearson: *r* = 0.17, *p* = 0.006). The *F. vesca* phenotype, in turn, was directly affected by mowing and fertilizer (Fig. 2A, B), the morphological and physiological features of plant communities (Fig. 2B), and also by microbiome composition (Fig. 2A) and diversity (Fig. 2B).

**Figure 2.**
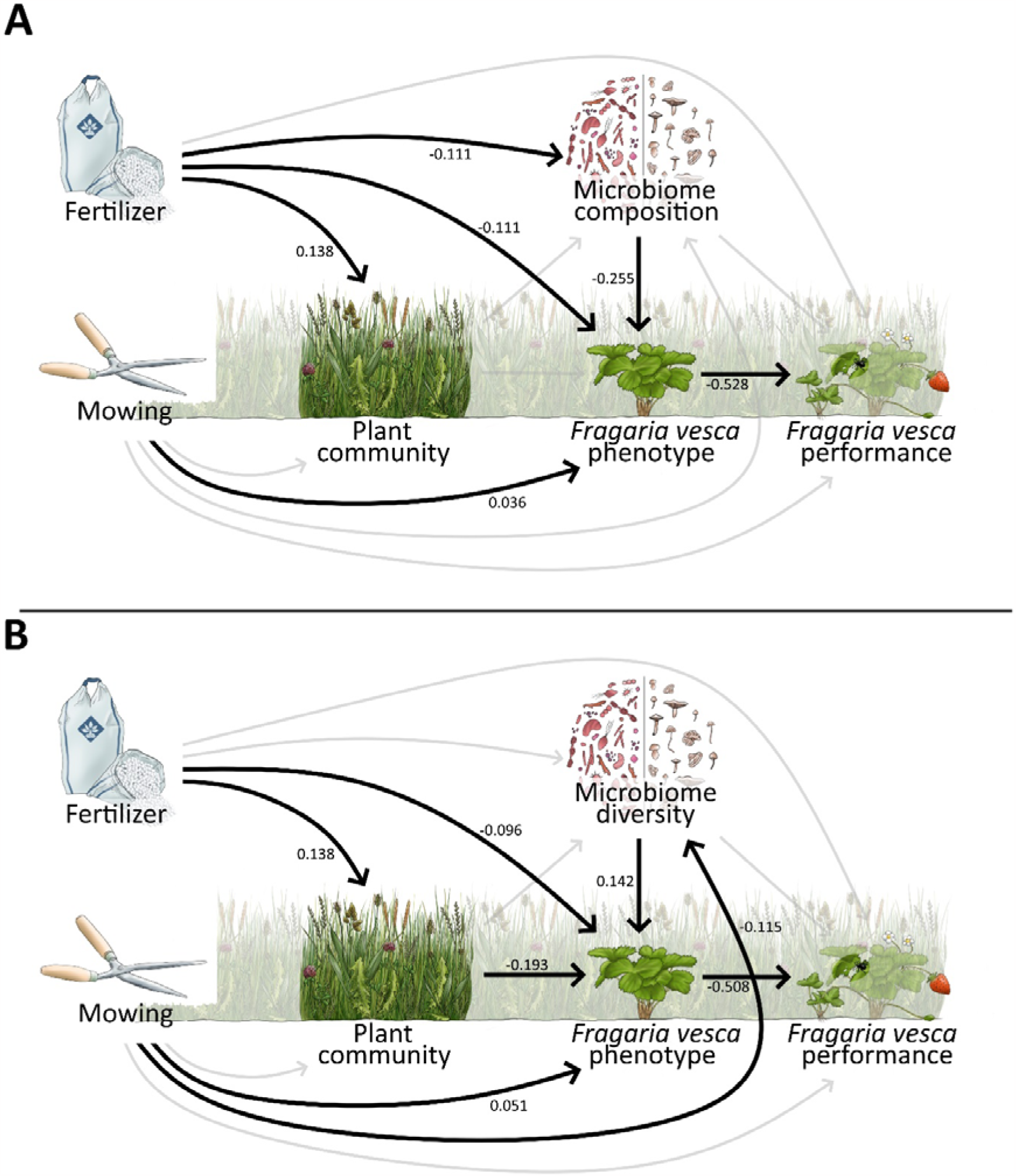
Effects of land-use components and microbiome on the phenotype and performance of *Fragaria vesca* phytometers in a common garden experiment. Structural equation models (SEM) testing effects of land-use components, plant communities, and plant microbiome on the phenotype and performance of individual plants. One SEM is considering the microbiome composition (A), the other one the microbiome diversity (B). Numbers next to arrows represent standardized path coefficients and are given for significant paths only (bold arrows). Statistical results are reported in the text and Supplementary Material (Tables 3).

In accordance with our second hypothesis, estimates of microbiome effects on *F. vesca* phenotype were stronger than those of the surrounding plant community and land-use components (Fig. 2A, B). Particularly some physiological traits were individually related to differences in microbiome composition (Pearson correlation; greenness: *p* = 0.049, R = 0.15; hue: *p* = 0.015, R = 0.19; NDVI: *p* = 0.020, R = 0.18). The multivariate *F. vesca* phenotype also corresponded to the multivariate representation of plant community features (Mantel: *r* = 0.15, *p* = 0.002). Some morphological and physiological traits of *F. vesca* and features of plant communities were directly correlated, too (Pearson correlation; leaf area: *p* = 0.002, *R* = 0.24; leaf area index: *p* < 0.001, *R* = 0.26; leaf area projected: *p* = 0.026, *R* = 0.18; leaf inclination: *p* = 0.003, *R* = -0.23; NPCI: *p* = 0.022, *R* = 0.18). Morphological and physiological features of plant communities as well as microbiome composition were affected by fertilizer addition (Fig. 2A, B), whereas microbiome diversity was affected by mowing treatments (Fig. 2B). Both structural equation models had a very good model fit (see Supplementary Table 3). Additional results supporting the effects shown in structural equation models can be found in Supplementary Tables 4 to 8. In summary, these results verify the hypotheses that the plant microbiome has stronger effects on the plant phenotype (and in consequence on plant performance) than direct land-use effects, and that plant performance is not a direct function of land-use intensity, supporting hypotheses two and three.

### Microbial effects on plant phenotype in the absence of other biotic or abiotic influences

To test our fourth hypothesis that microbial effects are also measurable in the absence of other biotic of abiotic influences, we performed an additional lab experiment. We cultivated *F. vesca* plants from surface-sterilized seeds in germ-free containers and inoculated these plants with the whole microbiomes of the *F. vesca* phytometer plants retrieved from the field experiment. Thus, inoculated plants differed only in microbiome composition and diversity, but shared identical environmental condition. Inoculated plants were phenotyped three months after the inoculation using the plant scanner introduced above and the parameters describing morphological and physiological traits of *F. vesca* plants were used as dependent variable in the following statistical analysis. The multivariate phenotype of *F. vesca* clearly responded to microbiome origin, i.e. land-use effects on the microbiome of donor plants (field-grown phytometers) were visible in the phenotype of lab-grown plants that did not differ in any abiotic treatments but in microbiome composition and diversity (CAP based on Bray–Curtis distances; mowing: *F*_1,91_ = 0.75, *p* = 0.68; fertilizer: *F*_1,91_ = 0.52, *p* = 0.96; mowing x fertilizer: *F*_1,91_ = 1.99, *p* = 0.028; Fig. 3A).

**Figure 3.**
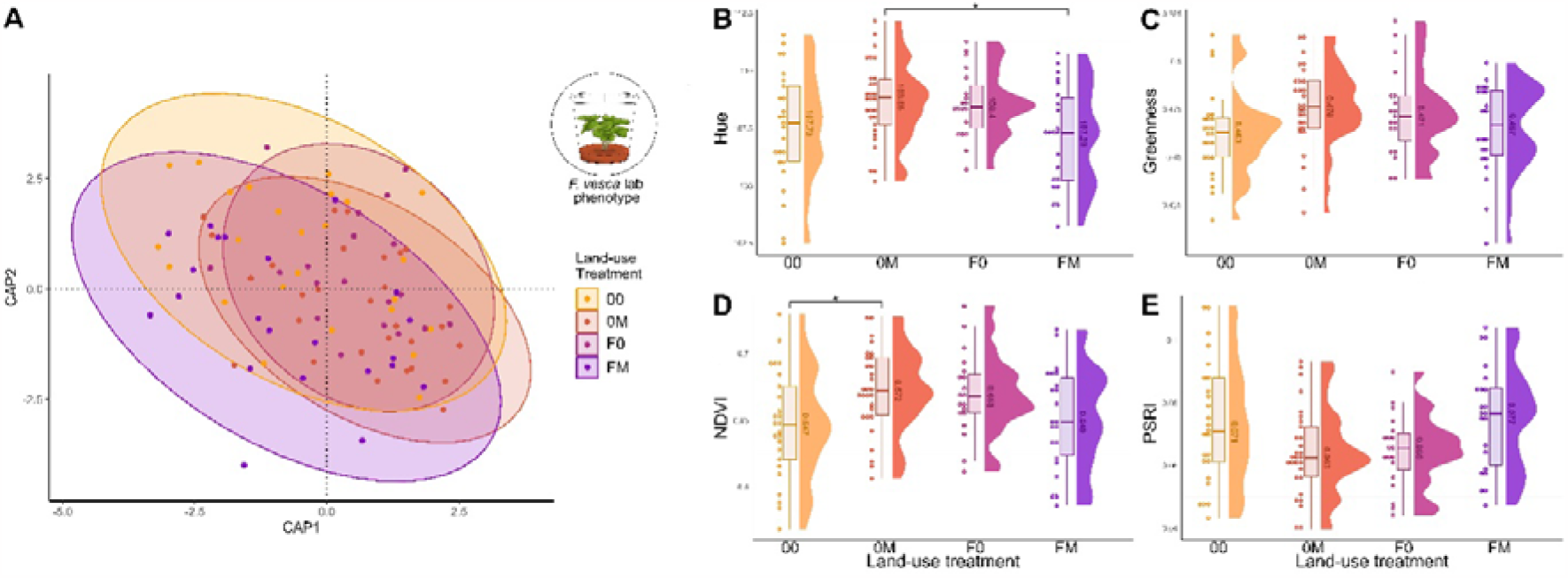
Phenotype of *F. vesca* plants inoculated with the microbiome of field-grown phytometers that were exposed to different land-use treatments. *F. vesca* plants grew from surface-sterilized seeds in germ-free containers and experienced identical conditions until phenotyping; they only differed in their microbiome. Microbiome-mediated land-use treatment effects on (A) the multivariate *F. vesca* phenotype or (B-E) individual physiological plant traits (Hue, Greenness, **N**ormalized **D**ifference **V**egetation **I**ndex, **P**lant **S**enescence **R**eflectance Index). Each circle represents one sample, land-use treatment is color-coded, significance is indicated with asterisks. Statistical results are reported in the text and Supplementary Material (Table 8).

Interestingly, some physiological traits of lab-grown plants individually responded to land-use treatments (ANOVA of linear model with mowing x fertilizer; hue: *F*_1,91_ = 9.4, *p* = 0.003; greenness: *F*_1,91_ = 5.9, *p* = 0.017; NDVI: *F*_1,91_ = 8.9, *p* = 0.004; PSRI: *F*_1,91_ = 8.9, *p* = 0.004, Fig. 3B-E), which corresponds to those traits that directly responded to the microbiome in field-grown phytometer plants (see above).

## Discussion

Our results largely verified our initial hypotheses on microbiome-assisted plant tolerance against land-use changes. The microbiome of the *F. vesca* phytometers rapidly altered composition and diversity in response to mowing and fertilization. These alterations in the plant-associated microbiome affected the phytometers phenotype – an effect that was stronger than direct land-use effects on morphological and physiological traits of *F. vesca*. As the performance of the plants was solely related to the plant phenotype and invariant across land-use treatments, we concluded that the plant microbiome mediates land-use effects on plant phenotype and performance and increases the tolerance of plants to environmental changes. This conclusion was clearly supported by our laboratory experiments that demonstrate the effect of the microbiome on the plant phenotype in the absence of other biotic or abiotic influences associated with changes in land-use. In contrast to previous studies that transplanted microbiomes from sites with different levels of environmental stress to increase plant tolerance (Allsup et al., 2023; Carrell et al., 2022; Kiesewetter & Afkhami, 2021; Porter et al., 2020; Timmusk et al., 2014), our data indicate the stress-mitigating potential of rapid modulations of the plant microbiome occurring on site. These natural modulations of the plant microbiome can be the key in plant tolerance to global change components and thus a potential pathway in addition to ‘adapt or migrate’ strategies (Scheffers et al., 2016) for species persistence and ecosystem stability.

Land-use intensification poses a major threat to many ecosystems, leading to species loss and alterations in community structure, which can disrupt ecosystem functioning (Allan et al., 2015; Díaz et al., 2019). Relocation as a strategy to avoid environmental changes might often fail because of time constraints, continued habitat shrinking, or a lacking spatial connectivity within a species’ range of dispersal (Asamoah et al., 2021; Corlett & Westcott, 2013; Gómez-Ruiz & Lacher Jr, 2019; Renton et al., 2013); likewise, the process of evolutionary adaptation to rapid alterations also takes several generations and might be too slow in most cases (Etterson & Shaw, 2001). Our data experimentally support recent findings that land-use not only affects plants and plant communities, but also the diversity and composition of plant-associated bacterial and fungal communities (Bai et al., 2023; Estendorfer et al., 2017; Gaube et al., 2021). Our data additionally demonstrate the pace and small scale of such alterations: Within three months of exposure to different land-use treatments in the small area of the common garden, surrounded by the same environment, microbial communities clearly diverged to treatment-specific compositions. Thus, our data support the idea that alterations in the microbiome can easily keep pace with changes in environmental conditions (Martiny et al., 2023). These rapid alterations are the prerequisite for microbiome-assisted plant tolerance to changing environmental conditions, which has been proposed in transplantation studies (Afkhami, 2023; Allsup et al., 2023). Our lab experiment provides important evidence – lacking in many other studies – of the contribution of the microbiome to the plant phenotype, which translates, according to our data, to plant performance. Therefore, small-scale natural variation in the plant microbiome in response to environmental conditions may be an untapped resource for microbiome engineering (Afridi et al., 2022; Rodriguez & Duran, 2020) that aids ecosystem restoration (Doering et al., 2021).

Microorganisms play essential roles in ecosystem tolerance and resilience against global change components (Hacquard et al., 2022; Saikkonen et al., 2020; Sessitsch et al., 2023). Here we demonstrated that rapid alterations of the plant microbiome may be key in plant species’ persistence despite changing environments and thus in ecosystem stability. However, a prerequisite for this microbiome-mediated strategy is locally available diverse sources of microorganisms that provide the source for rapid modulations in response to change. The need for a diverse selection of microbes is further supported by the finding that the strongest effects on plants are usually mediated by microbial consortia, not individual strains (Liu et al., 2023), and that microbial diversity can enhance plant species richness and productivity (Abrahao et al., 2022; Laforest-Lapointe et al., 2017). Therefore, we conclude that conservation efforts will benefit from considering and protecting microbial diversity in addition to efforts to maintain plant and animal communities. Given the multiple beneficial roles of microbes in natural and anthropogenically altered environments, such ‘microbiome stewardship’ (Peixoto et al., 2022) can help mitigate the effects of global change and facilitate environmental and human health.

## Acknowledgements

We thank the Botanical Garden of the University of Marburg for their support in establishing and maintaining the common garden and Annette Schriever and Alexander Lach for technical support. We thank the managers of the three Exploratories, Julia Bass, Anna K. Franke, Miriam Teuscher and all former managers for their work in maintaining the plot and project infrastructure; Christiane Fischer and Victoria Grießmeier for giving support through the central office, Andreas Ostrowski for managing the central data base, and Markus Fischer, Eduard Linsenmair, Dominik Hessenmöller, Daniel Prati, Ingo Schöning, François Buscot, Ernst-Detlef Schulze, Wolfgang W. Weisser and the late Elisabeth Kalko for their role in setting up the Biodiversity Exploratories project. We thank the administration of the Hainich national park, the UNESCO Biosphere Reserve Swabian Alb and the UNESCO Biosphere Reserve Schorfheide-Chorin as well as all land owners for the excellent collaboration. The work has been (partly) funded by the DFG Priority Program 1374 ‘Biodiversity Exploratories’ (JU 2856/4-1). Field work permits were issued by the responsible state environmental offices of Baden-Württemberg, Thüringen, and Brandenburg.

## Author contributions

RRJ conceived the study, VZ and RRJ designed the study, VZ performed field and lab work, AM provided material, VZ and RRJ performed statistical analysis, VZ and RRJ drafted the first version of the manuscript, all authors contributed to the final version.

## Data availability

This work was performed within the scope and using resources of the project EXClAvE of the Biodiversity Exploratories program (DFG Priority Program 1374). The dataset generated and analyzed during the current study is publicly available in the Biodiversity Exploratories Information System (id 31497, http://doi.org/10.17616/R32P9Q). The raw sequences of next-generation 16S and ITS amplicon sequencing are available at the European Nucleotide Archive ENA under the project accession PRJEB52951 (https://www.ebi.ac.uk/ena/browser/view/PRJEB52951).

## References

Abrahao, A., Marhan, S., Boeddinghaus, R. S., Nawaz, A., Wubet, T., Holzel, N., Klaus, V. H., Kleinebecker, T., Freitag, M., Hamer, U., Oliveira, R. S., Lambers, H., & Kandeler, E. (2022, Dec). Microbial drivers of plant richness and productivity in a grassland restoration experiment along a gradient of land-use intensity. New Phytol, 236(5), 1936–1950. 10.1111/nph.18503

Afkhami, M. E. (2023). Past microbial stress benefits tree resilience. Sci, 380.6647, 798–799. 10.1126/science.adi1594

Afridi, M. S., Javed, M. A., Ali, S., De Medeiros, F. H. V., Ali, B., Salam, A., Sumaira Marc, R. A., Alkhalifah, D. H. M., Selim, S., & Santoyo, G. (2022). New opportunities in plant microbiome engineering for increasing agricultural sustainability under stressful conditions. Front Plant Sci, 13, 899464. 10.3389/fpls.2022.899464

Allan, E., Manning, P., Alt, F., Binkenstein, J., Blaser, S., Bluthgen, N., Bohm, S., Grassein, F., Holzel, N., Klaus, V. H., Kleinebecker, T., Morris, E. K., Oelmann, Y., Prati, D., Renner, S. C., Rillig, M. C., Schaefer, M., Schloter, M., Schmitt, B., Schoning, I., Schrumpf, M., Solly, E., Sorkau, E., Steckel, J., Steffen-Dewenter, I., Stempfhuber, B., Tschapka, M., Weiner, C. N., Weisser, W. W., Werner, M., Westphal, C., Wilcke, W., & Fischer, M. (2015, Aug). Land use intensification alters ecosystem multifunctionality via loss of biodiversity and changes to functional composition. Ecol Lett, 18(8), 834–843. 10.1111/ele.12469

Allsup, C. M., George, I., & Lankau, R. A. (2023). Shifting microbial communities can enhance tree tolerance to changing climates. Sci, 380.6647, 835–840. 10.1126/science.adf2027

Amir, A., McDonald, D., Navas-Molina, J. A., Kopylova, E., Morton, J. T., Zech Xu, Z., Kightley, E. P., Thompson, L. R., Hyde, E. R., Gonzalez, A., & Knight, R. (2017, Mar-Apr). Deblur Rapidly Resolves Single-Nucleotide Community Sequence Patterns. mSystems, 2(2). 10.1128/mSystems.00191-16

Angevine, M. W. (1983). Variations in the demography of natural populations of the wild strawberries Fragaria vesca and F. virginiana. J Ecol, 959–974. 10.2307/2259605

Asamoah, E. F., Beaumont, L. J., & Maina, J. M. (2021). Climate and land-use changes reduce the benefits of terrestrial protected areas. Nat Clim Chang, 11(12), 1105–1110. 10.1038/s41558-021-01223-2

Bai, R., Hu, H. W., Zhou, M., Sheng, J., Xiong, C., Guo, Y., Yuan, Y., Hou, L., Zhang, W. H., & Bai, W. (2023). Effects of long-term mowing on leaf- and root-associated bacterial community structures are linked to functional traits in 11 plant species from a temperate steppe. Funct Ecol, 37(7), 1787–1801. 10.1111/1365-2435.14343

Bivand, R. (2022). R Packages for Analyzing Spatial Data: A Comparative Case Study with Areal Data. Geogr Anal, 54(3), 488–518. 10.1111/gean.12319

Bivand, R. S., Pebesma, E., & Gómez-Rubio, V. (2013). Spatial data import and export. Applied spatial data analysis with R, 83–125.

Blüthgen, N., Dormann, C. F., Prati, D., Klaus, V. H., Kleinebecker, T., Hölzel, N., Alt, F., Boch, S., Gockel, S., Hemp, A., Müller, J., Nieschulze, J., Renner, S. C., Schöning, I., Schumacher, U., Socher, S. A., Wells, K., Birkhofer, K., Buscot, F., Oelmann, Y., Rothenwöhrer, C., Scherber, C., Tscharntke, T., Weiner, C. N., Fischer, M., Kalko, E. K. V., Linsenmair, K. E., Schulze, E.-D., & Weisser, W. W. (2012). A quantitative index of land-use intensity in grasslands: integrating mowing, grazing and fertilization. Basic Appl Ecol, 13(3), 207–220. 10.1016/j.baae.2012.04.001

Bokulich, N. A., Kaehler, B. D., Rideout, J. R., Dillon, M., Bolyen, E., Knight, R., Huttley, G. A., & Gregory Caporaso, J. (2018). Optimizing taxonomic classification of marker-gene amplicon sequences with QIIME 2’s q2-feature-classifier plugin. Microbiome, 6(1), 1–17.

Bolyen, E., Rideout, J. R., Dillon, M. R., Bokulich, N. A., Abnet, C. C., Al-Ghalith, G. A., Alexander, H., Alm, E. J., Arumugam, M., & Asnicar, F. (2019). Reproducible, interactive, scalable and extensible microbiome data science using QIIME 2. Nature Biotechnology, 37(8), 852–857.

Callahan, B. J., McMurdie, P. J., Rosen, M. J., Han, A. W., Johnson, A. J., & Holmes, S. P. (2016, Jul). DADA2: High-resolution sample inference from Illumina amplicon data. Nat Methods, 13(7), 581–583. 10.1038/nmeth.3869

Carrell, A. A., Lawrence, T. J., Cabugao, K. G. M., Carper, D. L., Pelletier, D. A., Lee, J. H., Jawdy, S. S., Grimwood, J., Schmutz, J., Hanson, P. J., Shaw, A. J., & Weston, D. J. (2022, Jun). Habitatadapted microbial communities mediate Sphagnum peatmoss resilience to warming. New Phytol, 234(6), 2111–2125. 10.1111/nph.18072

Corlett, R. T., & Westcott, D. A. (2013). Will plant movements keep up with climate change? Trends Ecol Evol, 28(8), 482–488. 10.1016/j.tree.2013.04.003

Díaz, S., Settele, J. ES. B., Ngo, H. T., Guèze, M., Agard, J., Arneth, A., Balvanera, P., Brauman, K. A., Butchart, S. H. M., Chan, K. M. A., Garibaldi, L. A., Ichii, K., Liu, J., Subramanian, S. M., Midgley, G. F., Miloslavich, P., Molnár, Z., Obura, D., Pfaff, A., Polasky, S., Purvis, A., Razzaque, J., Reyers, B., Roy Chowdhury, R., Shin, Y. J., Visseren-Hamakers, I. J., Willis, K. J., & N., Z. C. (2019). IPBES (2019): Summary for policymakers of the global assessment report on biodiversity and ecosystem services of the Intergovernmental Science-Policy Platform on Biodiversity and Ecosystem Services. IPBES secretariat, Bonn, Germany, 56 pages.

Doering, T., Wall, M., Putchim, L., Rattanawongwan, T., Schroeder, R., Hentschel, U., & Roik, A. (2021). Towards enhancing coral heat tolerance: a “microbiome transplantation” treatment using inoculations of homogenized coral tissues. Microbiome, 9(1), 102. 10.1186/s40168-021-01053-6

Egan, P. A., Muola, A., & Stenberg, J. A. (2018). Capturing genetic variation in crop wild relatives: An evolutionary approach. Evol Appl, 11(8), 1293–1304. 10.1111/eva.12626

Engelsdorf, T., Gigli-Bisceglia, N., Veerabagu, M. M. J. F. Vaahtera, L., Augstein, F., Van der Does, D., Zipfel, C., & Hamann, T. (2018). The plant cell wall integrity maintenance and immune signaling systems cooperate to control stress responses in Arabidopsis thaliana. Sci Signal, 11(536), eaao3070. 10.1126/scisignal.aao3070

Estendorfer, J., Stempfhuber, B., Haury, P., Vestergaard, G., Rillig, M. C., Joshi, J., Schroder, P., & Schloter, M. (2017). The Influence of Land Use Intensity on the Plant-Associated Microbiome of Dactylis glomerata L. Front Plant Sci, 8, 930. 10.3389/fpls.2017.00930

Etterson, J. R., & Shaw, R. G. (2001). Constraint to adaptive evolution in response to global warming. Sci, 294.5540, 151–154. 10.1126/science.1063656

Feeley, K. J., Rehm, E. M., & Machovina, B. (2012). Perspective: the responses of tropical forest species to global climate change: acclimate, adapt, migrate, or go extinct? Front Biogeogr, 4(2). 10.21425/F5FBG12621

Finkel, O. M., Castrillo, G., Herrera Paredes, S., Salas Gonzalez, I., & Dangl, J. L. (2017, Aug). Understanding and exploiting plant beneficial microbes. Curr Opin Plant Biol, 38, 155–163. 10.1016/j.pbi.2017.04.018

Fischer, M., Bossdorf, O., Gockel, S., Hänsel, F., Hemp, A., Hessenmöller, D., Korte, G., Nieschulze, J., Pfeiffer, S., Prati, D., Renner, S., Schöning, I., Schumacher, U., Wells, K., Buscot, F., Kalko, E. K. V., Linsenmair, K. E., Schulze, E.-D., & Weisser, W. W. (2010). Implementing large-scale and long-term functional biodiversity research: The Biodiversity Exploratories. Basic Appl Ecol, 11(6), 473–485. 10.1016/j.baae.2010.07.009

Friesen, M. L., Porter, S. S., Stark, S. C., von Wettberg, E. J., Sachs, J. L., & Martinez-Romero, E. (2011). Microbially Mediated Plant Functional Traits. Annu Rev Ecol Evol Syst, 42(1), 23–46. 10.1146/annurev-ecolsys-102710-145039

Gaube, P., Junker, R. R., & Keller, A. (2021). Changes amid constancy: Flower and leaf microbiomes along land use gradients and between bioregions. Basic Appl Ecol, 50, 1–15. 10.1016/j.baae.2020.10.003

Gómez-Ruiz, E. P., & Lacher Jr, T. E. (2019). Climate change, range shifts, and the disruption of a pollinator-plant complex. Sci Rep, 9(1), 14048. 10.1038/s41598-019-50059-6

Gonzalez, A., Navas-Molina, J. A., Kosciolek, T., McDonald, D., Vazquez-Baeza, Y., Ackermann, G., DeReus, J., Janssen, S., Swafford, A. D., Orchanian, S. B., Sanders, J. G., Shorenstein, J., Holste, H., Petrus, S., Robbins-Pianka, A., Brislawn, C. J., Wang, M., Rideout, J. R., Bolyen, E., Dillon, M., Caporaso, J. G., Dorrestein, P. C., & Knight, R. (2018, Oct). Qiita: rapid, web-enabled microbiome meta-analysis. Nat Methods, 15(10), 796–798. 10.1038/s41592-018-0141-9

Gossner, M. M., Lewinsohn, T. M., Kahl, T., Grassein, F., Boch, S., Prati, D., Birkhofer, K., Renner, S. C., Sikorski, J., Wubet, T., Arndt, H., Baumgartner, V., Blaser, S., Bluthgen, N., Borschig, C., Buscot, F., Diekotter, T., Jorge, L. R., Jung, K., Keyel, A. C., Klein, A. M., Klemmer, S., Krauss, J., Lange, M., Muller, J., Overmann, J., Pasalic, E., Penone, C., Perovic, D. J., Purschke, O., Schall, P., Socher, S. A., Sonnemann, I., Tschapka, M., Tscharntke, T., Turke, M., Venter, P. C., Weiner, C. N., Werner, M., Wolters, V., Wurst, S., Westphal, C., Fischer, M., Weisser, W. W., & Allan, E. (2016, Dec 8). Land-use intensification causes multitrophic homogenization of grassland communities. Nature, 540(7632), 266–269. 10.1038/nature20575

Grace, J. B. (2006). Structural Equation Modeling and Natural Systems. Cambridge University Press. Cambridge, UK. 10.1017/CBO9780511617799

Grace, J. B., & Bollen, K. A. (2005). Interpreting the Results from Multiple Regression and Structural Equation Models. Bulletin of the Ecological Society of America, 86, 283–295.

Grigulis, K., Lavorel, S., Krainer, U., Legay, N., Baxendale, C., Dumont, M., Kastl, E., Arnoldi, C., Bardgett, R. D., Poly, F., Pommier, T., Schloter, M., Tappeiner, U., Bahn, M., Clément, J.-C., & Hutchings, M. (2013). Relative contributions of plant traits and soil microbial properties to mountain grassland ecosystem services. J Ecol, 101(1), 47–57. 10.1111/1365-2745.12014

Hacquard, S., Wang, E., Slater, H., & Martin, F. (2022). Impact of global change on the plant microbiome. New Phytol, 234(6), 1907–1909.

Hanusch, M., He, X., Ruiz-Hernandez, V., & Junker, R. R. (2022, May 6). Succession comprises a sequence of threshold-induced community assembly processes towards multidiversity. Commun Biol, 5(1), 424. 10.1038/s42003-022-03372-2

Hilmarsson, H. S., Hytönen, T., Isobe, S., Göransson, M., Toivainen, T., & Hallsson, J. H. (2017). Population genetic analysis of a global collection of Fragaria vesca using microsatellite markers. PLoS One, 12(8), e0183384. 10.1371/journal.pone.0183384

Jiang, M., Delgado-Baquerizo, M., Yuan, M. M., Ding, J., Yergeau, E., Zhou, J., Crowther, T. W., & Liang, Y. (2023, Jul). Home-based microbial solution to boost crop growth in low-fertility soil. New Phytol, 239(2), 752–765. 10.1111/nph.18943

Junker, R. R., He, X., Otto, J. C., Ruiz-Hernandez, V., & Hanusch, M. (2021, Sep 28). Divergent assembly processes? A comparison of the plant and soil microbiome with plant communities in a glacier forefield. FEMS Microbiol Ecol, 97(10). 10.1093/femsec/fiab135

Kaiser, K., Wemheuer, B., Korolkow, V., Wemheuer, F., Nacke, H., Schoning, I., Schrumpf, M., & Daniel, R. (2016, Sep 21). Driving forces of soil bacterial community structure, diversity, and function in temperate grasslands and forests. Sci Rep, 6, 33696. 10.1038/srep33696

Kiesewetter, K. N., & Afkhami, M. E. (2021, Nov). Microbiome-mediated effects of habitat fragmentation on native plant performance. New Phytol, 232(4), 1823–1838. 10.1111/nph.17595

Kim, Y. C., Leveau, J., McSpadden Gardener, B. B., Pierson, E. A., Pierson, L. S., 3rd, & Ryu, C. M. (2011, Mar). The multifactorial basis for plant health promotion by plant-associated bacteria. Appl Environ Microbiol, 77(5), 1548–1555. 10.1128/AEM.01867-10

Kõljalg, U., Nilsson, H. R., Schigel, D., Tedersoo, L., Larsson, K.-H., May, T. W., Taylor, A. F., Jeppesen, T. S., Frøslev, T. G., & Lindahl, B. D. (2020). The taxon hypothesis paradigm—on the unambiguous detection and communication of taxa. Microorganisms, 8(12), 1910.

Laforest-Lapointe, I., Paquette, A., Messier, C., & Kembel, S. W. (2017, Jun 1). Leaf bacterial diversity mediates plant diversity and ecosystem function relationships. Nature, 546(7656), 145–147. 10.1038/nature22399

Lau, J. A., & Lennon, J. T. (2012, Aug 28). Rapid responses of soil microorganisms improve plant fitness in novel environments. Proc Natl Acad Sci U S A, 109(35), 14058–14062. 10.1073/pnas.1202319109

Li, X., Jousset, A., de Boer, W., Carrion, V. J., Zhang, T., Wang, X., & Kuramae, E. E. (2019, Mar). Legacy of land use history determines reprogramming of plant physiology by soil microbiome. ISME J, 13(3), 738–751. 10.1038/s41396-018-0300-0

Liu, X., Mei, S., & Salles, J. F. (2023). Inoculated microbial consortia perform better than single strains in living soil: A meta-analysis. Appl Soil Ecol, 190. 10.1016/j.apsoil.2023.105011

Love, M. I., Huber, W., & Anders, S. (2014). Moderated estimation of fold change and dispersion for RNA-seq data with DESeq2. Genome Biol, 15(12), 1–21. 10.1186/s13059-014-0550-8

Manning, P., Gossner, M. M., Bossdorf, O., Allan, E., Zhang, Y.-Y., Prati, D., Blüthgen, N., Boch, S., Böhm, S., Börschig, C., Hölzel, N., Jung, K., Klaus, V. H., Klein, A. M. K. T., Krauss, J., Lange, M., Müller, J., Pašalić, E., Socher, S. A., Tschapka, M., Türke, M., Weiner, C., Werner, M., Gockel, S., Hemp, A., Renner, S. C., Wells, K., Buscot, F., Kalko, E. K. V., Linsenmair, K. E., Weisser, W. W., & Fischer, M. (2015). Grassland management intensification weakens the associations among the diversities of multiple plant and animal taxa. Ecology, 96(6), 1492–1501.

Martiny, J. B. H., Martiny, A. C., Brodie, E., Chase, A. B., Rodriguez-Verdugo, A., Treseder, K. K., & Allison, S. D. (2023, Mar 25). Investigating the eco-evolutionary response of microbiomes to environmental change. Ecol Lett. 10.1111/ele.14209

McDonald, D., Price, M. N., Goodrich, J., Nawrocki, E. P., DeSantis, T. Z., Probst, A., Andersen, G. L., Knight, R., & Hugenholtz, P. (2012). An improved Greengenes taxonomy with explicit ranks for ecological and evolutionary analyses of bacteria and archaea. The ISME journal, 6(3), 610–618.

Muola, A., Weber, D., Malm, L. E., Egan, P. A., Glinwood, R., Parachnowitsch, A. L., & Stenberg, J. A. (2017). Direct and pollinator-mediated effects of herbivory on strawberry and the potential for improved resistance. Front Plant Sci, 8, 823. 10.3389/fpls.2017.00823

Newbold, T., Hudson, L. N., Hill, S. L., Contu, S., Lysenko, I., Senior, R. A., Borger, L., Bennett, D. J., Choimes, A., Collen, B., Day, J., De Palma, A., Diaz, S., Echeverria-Londono, S., Edgar, M. J., Feldman, A., Garon, M., Harrison, M. L., Alhusseini, T., Ingram, D. J., Itescu, Y., Kattge, J., Kemp, V., Kirkpatrick, L., Kleyer, M., Correia, D. L., Martin, C. D., Meiri, S., Novosolov, M., Pan, Y., Phillips, H. R., Purves, D. W., Robinson, A., Simpson, J., Tuck, S. L., Weiher, E., White, H. J., Ewers, R. M., Mace, G. M., Scharlemann, J. P., & Purvis, A. (2015, Apr 2). Global effects of land use on local terrestrial biodiversity. Nature, 520(7545), 45–50. 10.1038/nature14324

Oksanen, J., Simpson, G., Blanchet, F., Kindt, R., Legendre, P., Minchin, P., O’Hara, R., Solymos, P., Stevens, M., Szoecs, E., Wagner, H., Barbour, M., Bedward, M., Bolker, B., Borcard, D., Carvalho, G., Chirico, M., De Caceres, M., Durand, S., Evangelista, H., FitzJohn, R., Friendly, M., Furneaux, B., Hannigan, G., Hill, M., Lahti, L., McGlinn, D., Ouellette, M., Ribeiro Cunha, E., Smith, T., Stier, A., Ter Braak, C., & Weedon, J. (2022). vegan: Community Ecology Package. R package version 2.6-4. https://CRAN.R-project.org/package=vegan. In

Paulson, J. (2014). metagenomeSeq: statistical analysis for sparse high-throughput sequencing.

Peixoto, R. S., Voolstra, C. R., Sweet, M., Duarte, C. M., Carvalho, S., Villela, H., Lunshof, J. E., Gram, L., Woodhams, D. C., Walter, J., Roik, A., Hentschel, U., Thurber, R. V., Daisley, B., Ushijima, B., Daffonchio, D., Costa, R., Keller-Costa, T., Bowman, J. S., Rosado, A. S., Reid, G., Mason, C. E., Walke, J. B., Thomas, T., & Berg, G. (2022, Nov). Harnessing the microbiome to prevent global biodiversity loss. Nat Microbiol, 7(11), 1726–1735. 10.1038/s41564-022-01173-1

Porter, S. S., Bantay, R., Friel, C. A., Garoutte, A., Gdanetz, K., Ibarreta, K., Moore, B. M., Shetty, P., Siler, E., Friesen, M. L., & Bennett, A. (2020). Beneficial microbes ameliorate abiotic and biotic sources of stress on plants. Funct Ecol, 34(10), 2075–2086. 10.1111/1365-2435.13499

R Core Team. (2022). R: A language and environment for statistical computing. R Foundation for Statistical Computing, Vienna, Austria. https://www.R-project.org/. In

Renton, M., Childs, S., Standish, R., & Shackelford, N. (2013). Plant migration and persistence under climate change in fragmented landscapes: does it depend on the key point of vulnerability within the lifecycle? Ecol Modell, 249, 50–58. 10.1016/j.ecolmodel.2012.07.005

Rodriguez, R., & Duran, P. (2020). Natural Holobiome Engineering by Using Native Extreme Microbiome to Counteract the Climate Change Effects. Front Bioeng Biotechnol, 8, 568. 10.3389/fbioe.2020.00568

Roiloa, S. R., & Retuerto, R. (2007). Responses of the clonal Fragaria vesca to microtopographic heterogeneity under different water and light conditions. Environ Exp Bot, 61(1), 1–9. 10.1016/j.envexpbot.2007.02.006

Rosseel, Y. (2012). lavaan: An R Package for Structural Equation Modeling. Journal of statistical software, 48, 1–36.

Saikkonen, K., Nissinen, R., & Helander, M. (2020). Toward Comprehensive Plant Microbiome Research. Front Ecol Evol, 8. 10.3389/fevo.2020.00061

Scheffers, B. R., De Meester, L., Bridge, T. C., Hoffmann, A. A., Pandolfi, J. M., Corlett, R. T., Butchart, S. H., Pearce-Kelly, P., Kovacs, K. M., Dudgeon, D., Pacifici, M., Rondinini, C., Foden, W. B., Martin, T. G., Mora, C., Bickford, D., & Watson, J. E. (2016, Nov 11). The broad footprint of climate change from genes to biomes to people. Sci, 354(6313). 10.1126/science.aaf7671

Schulze, J., Rufener, R., Erhardt, A., & Stoll, P. (2012). The relative importance of sexual and clonal reproduction for population growth in the perennial herb Fragaria vesca. Popul Ecol, 54, 369–380. 10.1007/s10144-012-0321-x

Sessitsch, A., Wakelin, S., Schloter, M., Maguin, E., Cernava, T., Champomier-Verges, M.-C., Charles, T. C., Cotter, P. D., Ferrocino, I., & Kriaa, A. (2023). Microbiome Interconnectedness throughout Environments with Major Consequences for Healthy People and a Healthy Planet. Microbiol Mol Biol Rev, e00212–00222. 10.1128/mmbr.00212-22

Socher, S. A., Prati, D., Boch, S., Müller, J., Baumbach, H., Gockel, S., Hemp, A., Schöning, I., Wells, K., Buscot, F., Kalko, E. K. V., Linsenmair, K. E., Schulze, E.-D., Weisser, W. W., & Fischer, M. (2013). Interacting effects of fertilization, mowing and grazing on plant species diversity of 1500 grasslands in Germany differ between regions. Basic Appl Ecol, 14(2), 126–136. 10.1016/j.baae.2012.12.003

Timmusk, S., Abd El-Daim, I. A., Copolovici, L., Tanilas, T., Kannaste, A., Behers, L., Nevo, E., Seisenbaeva, G., Stenstrom, E., & Niinemets, U. (2014). Drought-tolerance of wheat improved by rhizosphere bacteria from harsh environments: enhanced biomass production and reduced emissions of stress volatiles. PLoS One, 9(5), e96086. 10.1371/journal.pone.0096086

Trivedi, P., Batista, B. D., Bazany, K. E., & Singh, B. K. (2022, Jun). Plant-microbiome interactions under a changing world: responses, consequences and perspectives. New Phytol, 234(6), 1951–1959. 10.1111/nph.18016

Xirocostas, Z. A., Debono, S. A., Slavich, E., & Moles, A. T. (2022). The ZAX Herbivory Trainer—Free software for training researchers to visually estimate leaf damage. Methods Ecol Evol, 13(3), 596–602. 10.1111/2041-210X.13785

Yang, J., Kloepper, J. W., & Ryu, C. M. (2009, Jan). Rhizosphere bacteria help plants tolerate abiotic stress. Trends Plant Sci, 14(1), 1–4. 10.1016/j.tplants.2008.10.004

Zieschank, V., & Junker, R. R. (2023). Digital whole-community phenotyping: tracking morphological and physiological responses of plant communities to environmental changes in the field. Front Plant Sci, 14, 1141554. 10.3389/fpls.2023.1141554

