## Supplementary Material for "Tolerance to land-use changes through natural modulations of the plant microbiome"

**Supplementary Information**

**Supplementary Tables**

**Supplementary Table 1.** Land-use effects on bacterial and fungal diversity associated to leaves and roots of *Fragaria vesca* (grown in common garden). Results of ANOVA with mowing and fertilizer application explanatory variables are shown. Significant results are indicated in bold.

|  | sample | mowing | | fertilizer | |
| --- | --- | --- | --- | --- | --- |
|  |  | F_1,153_ | *p* | F_1,153_ | *p* |
| bacteria | leaf | 4.424 | **0.037** | 27.913 | **<0.001** |
|  | root | 0.868 | 0.353 | 2.038 | 0.155 |
| fungi | leaf | 9.501 | **0.002** | 0.785 | 0.377 |
|  | root | 1.890 | 0.171 | 4.046 | **0.046** |

**Supplementary Table 2.** Land-use effects on composition of bacterial and fungal communities associated to leaves and roots of *Fragaria vesca* (grown in common garden). Results of distance-based redundancy analyses using Bray-Curtis distances followed by a permutation test under reduced model with subsequent analysis of variance are shown. Significant results are indicated in bold.

|  | sample | mowing | | fertilizer | |
| --- | --- | --- | --- | --- | --- |
|  |  | F_1,153_ | *p* | F_1,153_ | *p* |
| bacteria | leaf | 1.733 | **<0.001** | 4.109 | **<0.001** |
|  | root | 1.299 | **0.014** | 3.156 | **<0.001** |
| fungi | leaf | 3.572 | **<0.001** | 2.937 | **<0.001** |
|  | root | 1.206 | **0.044** | 1.861 | **<0.001** |

**Supplementary Table 3.** Model fit parameters of both structural equation models (see Fig. 2A, B in the main text). SEM 1 uses microbiome composition, SEM 2 microbiome diversity as parameter.

| model | parameter | value |
| --- | --- | --- |
| SEM 1 (Fig. 2A) | *p*_chi-square_ | 0.771 |
|  | CFI | 1 |
|  | TLI | 1.108 |
|  | RMSEA | 0 |
|  | SRMR | 0.004 |
| SEM 2 (Fig. 2B) | *p*_chi-square_ | 0.666 |
|  | CFI | 1 |
|  | TLI | 1.102 |
|  | RMSEA | 0 |
|  | SRMR | 0.006 |

**Supplementary Table 4.** Land-use treatment (mowing and fertilizer application) effects on plant community, *F. vesca* phenotype (grown in common garden), microbiome composition, microbiome diversity, and *F. vesca* performance (grown in common garden). Results of distance-based redundancy analyses using Bray-Curtis distances followed by a permutation test under reduced model with subsequent analysis of variance (in the case of plant community, *F. vesca* phenotype, microbiome composition) or of ANOVA with mowing and fertilizer application explanatory variables (in the case of microbiome diversity, and *F. vesca* performance) are shown. Significant results are indicated in bold.

| plant community | F_1,158_ | *p* |
| --- | --- | --- |
| mowing | 7.87 | **<0.001** |
| fertilizer | 7.27 | **<0.001** |
| mowing x fertilizer | 196 | **0.030** |
| *F. vesca* phenotype |  |  |
| mowing | 1.15 | 0.271 |
| fertilizer | 10.53 | **<0.001** |
| mowing x fertilizer | 1.96 | **0.029** |
| microbiome composition |  |  |
| mowing | 1.69 | **<0.001** |
| fertilizer | 3.24 | **<0.001** |
| mowing x fertilizer | 1.08 | 0.157 |
| microbiome diversity |  |  |
| mowing | 8.05 | **0.005** |
| fertilizer | 1.87 | 0.173 |
| mowing x fertilizer | 0.49 | 0.483 |
| *F. vesca* performance |  |  |
| mowing | 1.98 | 0.290 |
| fertilizer | 2.51 | 0.120 |
| mowing x fertilizer | 0.06 | 0.811 |

**Supplementary Table 5.** Association between F*. vesca* (grown in the common garden) phenotype and performance or plant community features. Results of Mantel tests based on Pearson’s correlations (9999 permutations) calculated from Bray-Curtis distances are shown. Significant results are indicated in bold.

| matrix 1 | matrix 2 | r | *p* |
| --- | --- | --- | --- |
| *F. vesca* phenotype | *F. vesca* performance | 0.17 | **0.006** |
| *F. vesca* phenotype | plant community features | 0.15 | **0.002** |

**Supplementary Table 6.** Associations between microbiome composition (PC1) and plant community features (morphological and physiological parameters). Results of Pearson’s correlation are shown. Significant results are indicated in bold.

| parameter | microbiome composition | | plant community features | |
| --- | --- | --- | --- | --- |
|  | R | *p* | R | *p* |
| digital biomass | 0.15 | 0.054 | 0.08 | 0.300 |
| height | 0.15 | 0.061 | 0.11 | 0.150 |
| leaf angle | 0.12 | 0.120 | -0.14 | **0.027** |
| leaf area | 0.14 | 0.086 | 0.24 | **0.002** |
| leaf area index | 0.14 | 0.071 | 0.26 | **<0.001** |
| leaf area projected | 0.14 | 0.076 | 0.18 | **0.026** |
| leaf inclination | 0.02 | 0.830 | -0.23 | **0.003** |
| light penetration depth | 0.05 | 0.560 | 0.05 | 0.540 |
| greenness | 0.15 | **0.049** | -0.001 | 0.990 |
| hue | 0.19 | **0.015** | 0.08 | 0.330 |
| NDVI | 0.18 | **0.020** | 0.01 | 0.870 |
| NPCI | -0.05 | 0.490 | 0.18 | **0.022** |
| PSRI | -0.13 | 0.088 | 0.03 | 0.670 |

**Supplementary Table 7.** Land-use effects on morphological and physiological parameters of *F. vesca* phytometers grown in the common garden. Results of ANOVA with mowing and fertilizer as explanatory variables are shown. Significant results are indicated in bold.

| parameter | mowing | | fertilizer | | mowing x fertilizer | |
| --- | --- | --- | --- | --- | --- | --- |
|  | F_1,158_ | *p* | F_1,158_ | *p* | F_1,158_ | *p* |
| digital biomass | 3.860 | 0.051 | 45.020 | **<0.001** | 0.649 | 0.421 |
| height | 0.696 | 0.405 | 75.877 | **<0.001** | 0.901 | 0.344 |
| leaf angle | 0.832 | 0.363 | 6.393 | **0.012** | 0.001 | 0.975 |
| leaf area | 5.608 | **0.019** | 26.893 | **<0.001** | 2.687 | 0.103 |
| leaf area index | 4.087 | **0.045** | 23.951 | **<0.001** | 2.621 | 0.108 |
| leaf area projected | 4.195 | **0.042** | 16.438 | **<0.001** | 3.074 | 0.083 |
| leaf inclination | 1.824 | 0.179 | 21.330 | **<0.001** | 2.414 | 0.122 |
| light penetration depth | 0.414 | 0.521 | 19.683 | **<0.001** | 1.872 | 0.173 |
| greenness | 3.167 | 0.077 | 10.604 | **0.001** | 0.055 | 0.815 |
| hue | 0.031 | 0.860 | 0.110 | 0.741 | 0.521 | 0.472 |
| NDVI | 0.330 | 0.566 | 1.771 | 0.185 | 0.229 | 0.633 |
| NPCI | 1.251 | 0.265 | 1.474 | 0.227 | 1.606 | 0.207 |
| PSRI | 0.132 | 0.717 | 0.097 | 0.756 | 0.120 | 0.730 |

**Supplementary Table 8.** Microbiome-mediated effects on morphological and physiological parameters of *F. vesca* plants grown in lab under containment. Microbiomes originating from *F. vesca* plants growing in the common garden under different land-use treatments were inoculated on *F. vesca* plants in the lab. Results of ANOVA with mowing and fertilizer as explanatory variables are shown. Significant results are indicated in bold.

| parameter | mowing | | fertilizer | | mowing x fertilizer | |
| --- | --- | --- | --- | --- | --- | --- |
|  | F_1,91_ | *p* | F_1,91_ | *p* | F_1,91_ | *P* |
| digital biomass | 0.433 | 0.512 | 0.120 | 0.730 | 0.032 | 0.860 |
| height | 0.209 | 0.648 | 0.016 | 0.899 | 0.036 | 0.850 |
| leaf angle | 2.052 | 0.155 | 0.742 | 0.391 | 0.767 | 0.383 |
| leaf area | 0.362 | 0.549 | 0.196 | 0.659 | 0.004 | 0.951 |
| leaf area index | 0.270 | 0.604 | 0.221 | 0.639 | 0.004 | 0.950 |
| leaf area projected | 0.791 | 0.376 | 0.319 | 0.574 | 0.007 | 0.934 |
| leaf inclination | 1.527 | 0.220 | 0.840 | 0.362 | 0.713 | 0.401 |
| light penetration depth | 0.035 | 0.851 | 0.082 | 0.775 | 0.551 | 0.460 |
| greenness | 0.795 | 0.375 | 0.074 | 0.786 | 5.905 | **0.017** |
| hue | 0.004 | 0.952 | 0.690 | 0.409 | 9.443 | **0.003** |
| NDVI | 0.862 | 0.355 | 0.004 | 0.952 | 8.929 | **0.004** |
| NPCI | 0.047 | 0.830 | 0.171 | 0.680 | 2.787 | 0.098 |
| PSRI | 0.043 | 0.836 | 0.266 | 0.607 | 8.900 | **0.004** |

**Supplementary Table 9.** Moran’s I for endogenous variables of both structural equation models (see Fig. 2A,B in the main text) to test for spatial autocorrelation. Significant results are indicated in bold.

| model | variable | *p* |
| --- | --- | --- |
| SEM 1 (Fig. 2A) | plant community | <0.001 |
|  | *F. vesca* phenotype | 0.1715 |
|  | *F. vesca* performance | 0.07628 |
|  | microbiome composition | 0.2912 |
| SEM 2 (Fig. 2B) | plant community | <0.001 |
|  | *F. vesca* phenotype | 0.1534 |
|  | *F. vesca* performance | 0.08182 |
|  | microbiome diversity | 0.4728 |

**Supplementary Figures**

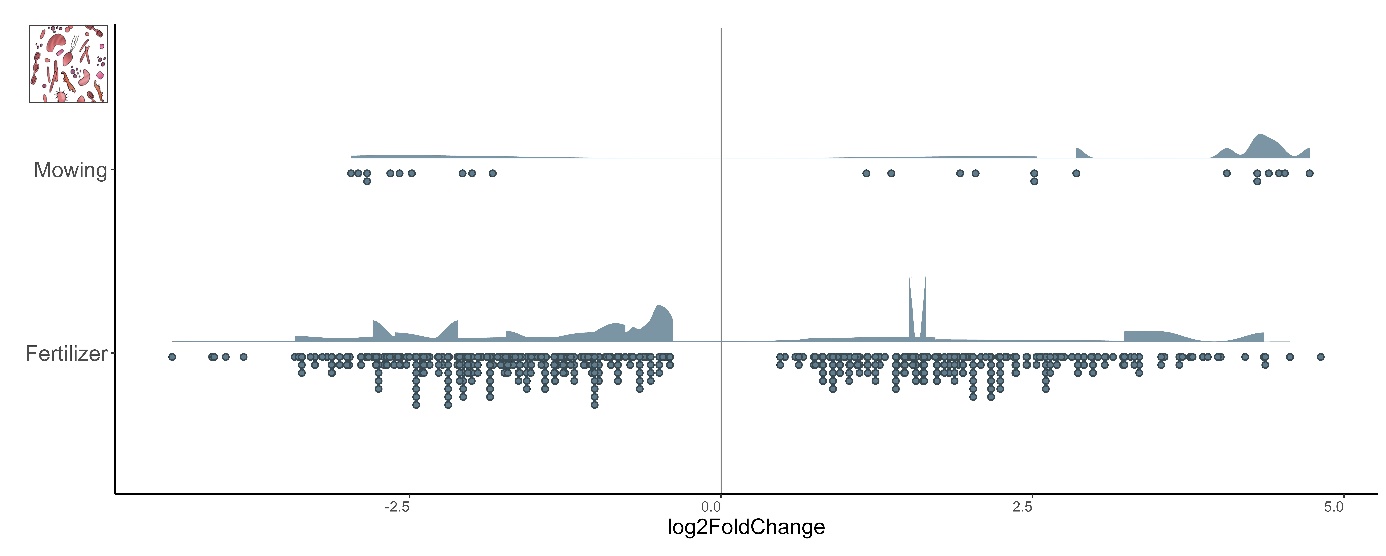

**Supplementary Figure 1.** Land-use effects on individual bacterial ASVs. ASVs that significantly responded to mowing or fertilizer application have been identified using the R package *DESeq2.* log2FoldChange gives an estimate for effect size.

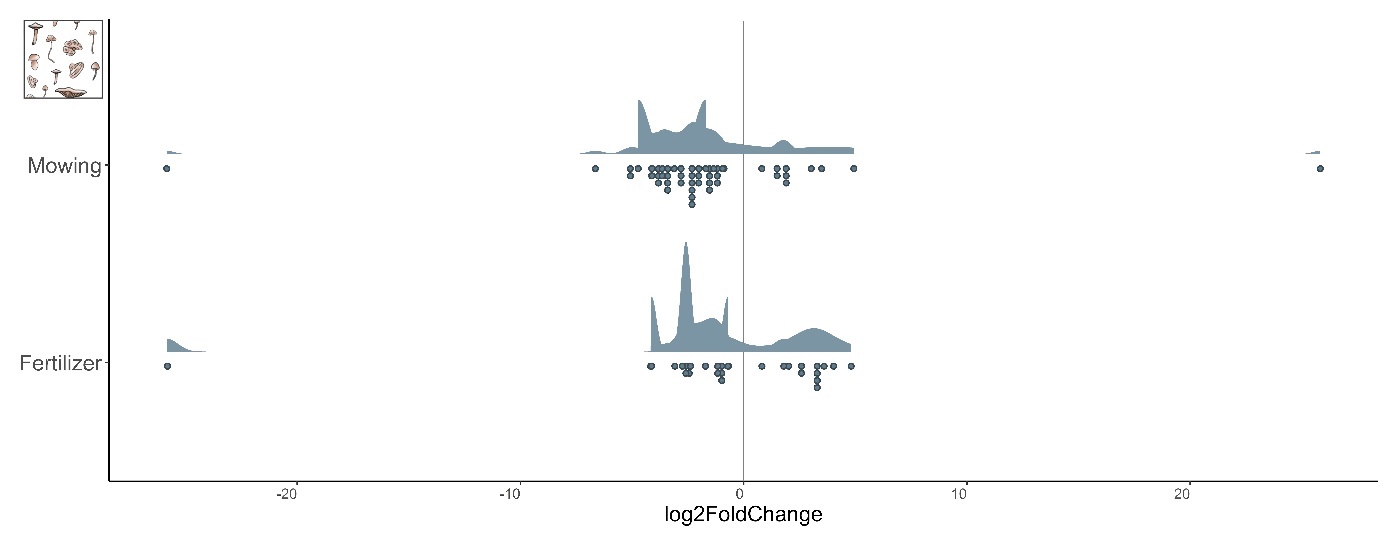

**Supplementary Figure 2.** Land-use effects on individual fungal ASVs. ASVs that significantly responded to mowing or fertilizer application have been identified using the R package *DESeq2.* log2FoldChange gives an estimate for effect size.
